## Supplementary References S1 S5 for "The macroevolutionary impact of recent and imminent mammal extinctions on Madagascar"

**References used to compile the checklist of Malagasy mammals (Data S1), the extinction data (Data S1) and the taxonomic state changes for recently up-listed species in the IUCN Red List (Data S5)**

1.	M. Kouvari, A. A. E. van der Geer, Biogeography of extinction: The demise of insular mammals from the Late Pleistocene till today. *Palaeogeogr. Palaeoclimatol. Palaeoecol.* **505**, 295–304 (2018).

2.	S. M. Goodman, H. H. Zafindranoro, V. Soarimalala, A case of the sympatric occurrence of *Microgale brevicaudata* and *M. grandidieri* (Afrosoricida, Tenrecidae) in the Beanka Forest, Maintirano. *Malagasy Nature.* **5**, 104-108 (2011).

3.	K. M. Everson, V. Soarimalala, S. M. Goodman, L. E. Olson, Multiple loci and complete taxonomic sampling resolve the phylogeny and biogeographic history of tenrecs (Mammalia: Tenrecidae) and reveal higher speciation rates in Madagascar’s humid forests. *Syst. Biol.* **65**, 890–909 (2016).

4.	S. M. Goodman, W. L. Jungers, *Extinct Madagascar: Picturing the Island’s Past* (University of Chicago Press, 2014).

15.	L. Werdelin, *Cenozoic Mammals of Africa* (University of California Press, 2010).

21.	S. M. Goodman, *Les Carnivora de Madagascar* (Association Vahatra in Antananarivo, 2012).

22.	B. D. Gerber, S. M. Karpanty, J. Randrianantenaina, Activity patterns of carnivores in the rain forests of Madagascar: implications for species coexistence. *J. Mammal.* **93**, 667–676 (2012).

23.	D. A. Burney, *et al.*, Subfossil lemur discoveries from the Beanka Protected Area in western Madagascar. *Quat. Res. U. S.* **93**, 187–203 (2020).

24.	S. M. Goodman, K. M. Helgen, Species limits and distribution of the Malagasy carnivoran genus *Eupleres* (Family Eupleridae). *Mammalia* **74**, 177–185 (2010).

25.	S. M. Goodman, *Les Chauves-Souris de Madagascar* (Association Vahatra in Antananarivo, 2012).

55.	R. Andriantompohavana, *Molecular phylogeny and taxonomic revision of the woolly lemurs, genus* Avahi *(Primates: Lemuriformes)* (Museum of Texas Tech University, 2007).

56.	J. P. Herrera, Testing the adaptive radiation hypothesis for the lemurs of Madagascar. *R. Soc. Open Sci.* **4**, 161014.

106.	R. M. Nowak, E. P. Walker, *Walker’s Mammals of the World* (JHU press, 1999).

107.	K. E. Samonds, Late Pleistocene bat fossils from Anjohibe Cave, northwestern Madagascar. *Acta Chiropterologica* **9**, 39–65 (2007).

108.	D. A. Burney, *et al.*, A chronology for late prehistoric Madagascar. *J. Hum. Evol.* **47**, 25–63 (2004).

109.	L. Godfrey, IUCN Red List of Threatened Species: *Palaeopropithecus* *ingens*. *IUCN Red List Threat. Species* (2021).

116.	J. R. Boisserie, IUCN Red List of Threatened Species: *Hippopotamus* *madagascariensis*. *IUCN Red List Threat. Species* (2016).

117.	S. M. Goodman, N. Vasey, D. A. Burney, Description of a new species of subfossil shrew tenrec (Afrosoricida: Tenrecidae: *Microgale*) from cave deposits in southeastern Madagascar. *Proc. Biol. Soc. Wash.* **120**, 367–376 (2007).

118.	S. T. Turvey, *Holocene Extinctions* (OUP Oxford, 2009).

119.	G. Veron, *et al.*, New insights into the systematics of Malagasy mongoose‐like carnivorans (Carnivora, Eupleridae, Galidiinae) based on mitochondrial and nuclear DNA sequences. *J. Zool. Syst. Evol. Res.* **55**, 250–264 (2017).

120.	R. A. Mittermeier, *et al.*, Lemur diversity in Madagascar. *Int. J. Primatol.* **29**, 1607–1656 (2008).

121.	P. Kappeler, Morphology, behaviour and molecular evolution of giant mouse lemurs (*Mirza* spp.) Gray, 1870, with description of a new species. *Primate Rep* **71**, 3–26 (2005).

122.	D. E. Wilson, D. M. Reeder, *Mammal Species of the World: A Taxonomic and Geographic Reference* (JHU Press, 2005).

123.	L. F. Groeneveld, D. W. Weisrock, R. M. Rasoloarison, A. D. Yoder, P. M. Kappeler, Species delimitation in lemurs: Multiple genetic loci reveal low levels of species diversity in the genus *Cheirogaleus*. *BMC Evol. Biol.* **9**, 1–16 (2009).

124.	International Union for Conservation of Nature, The IUCN Red List of Threatened Species. Version 2021-2. https://www.iucn.org/. Accessed November 10, 2021.

125.	International Union for Conservation of Nature, The IUCN Red List of Threatened Species. Version 2015-3. https://www.iucn.org/. Accessed November 10, 2021.

126.	International Union for Conservation of Nature, The IUCN Red List of Threatened Species. Version 2010.4. https://www.iucn.org/. Accessed November 10, 2021.
